# Biodiversity loss underlies the dilution effect of biodiversity

**DOI:** 10.1101/2020.04.20.050377

**Authors:** Fletcher W. Halliday, Jason R. Rohr, Anna-Liisa Laine

**Affiliations:** Department of Evolutionary Biology and Environmental Studies, University of Zurich, 8057, Zurich, CH; Department of Biological Sciences, Eck Institute of Global Health, Environmental Change Initiative, University of Notre Dame, Notre Dame, IN, USA; Organismal & Evolutionary Biology Research Program, PO Box 65, FI-00014 University of Helsinki, Finland

**Keywords:** biodiversity, parasitism, community structure, dilution effect

## Abstract

The dilution effect predicts increasing biodiversity to reduce the risk of infection, but the generality of this effect remains unresolved. Because biodiversity loss generates predictable changes in host community competence, we hypothesized that biodiversity loss might drive the dilution effect. We tested this hypothesis by reanalyzing four previously published meta-analyses that came to contradictory conclusions regarding generality of the dilution effect. In the context of biodiversity loss, our analyses revealed a unifying pattern: dilution effects were inconsistently observed for natural biodiversity gradients, but were commonly observed for biodiversity gradients generated by disturbances causing losses of native biodiversity. Incorporating biodiversity loss into tests of generality of the dilution effect further indicated that scale-dependency may strengthen the dilution effect only when biodiversity gradients are driven by biodiversity loss. Together, these results help to resolve one of the most contentious issues in disease ecology: the generality of the dilution effect.

## Introduction

Increasing biodiversity is often associated with a reduction in the risk of infectious diseases, a phenomenon known as the dilution effect (Keesing *et al.* 2006, 2010; Civitello *et al.* 2015; Halliday & Rohr 2019). Yet, despite more than three decades of empirical research, meta-analyses, reviews, and syntheses, there remains polarizing debate regarding the generality of this effect (Halsey 2019; Rohr *et al.* 2020). Several recent studies provide a promising framework for resolving this debate, suggesting that changes in the structure of host communities, rather than biodiversity per se, can explain when a dilution effect should be observed (Johnson *et al.* 2013, 2019; Joseph *et al.* 2013; Mihaljevic *et al.* 2014; Strauss *et al.* 2016; Liu *et al.* 2018; Halliday *et al.* 2019). Implicit in these studies is a focus on biodiversity loss: the structure of host communities often shifts predictably when biodiversity is lost or recovered, particularly following disturbances, and often in a way that favors species with combinations of physiological traits associated with increased disease risk (Joseph *et al.* 2013; Mihaljevic *et al.* 2014; Johnson *et al.* 2015a). These predictable shifts suggest that there should be a strong relationship between biodiversity and disease risk following a loss of native biodiversity. In contrast, such predictable changes are not expected over natural biodiversity gradients (Table 1).

**Table 1.**
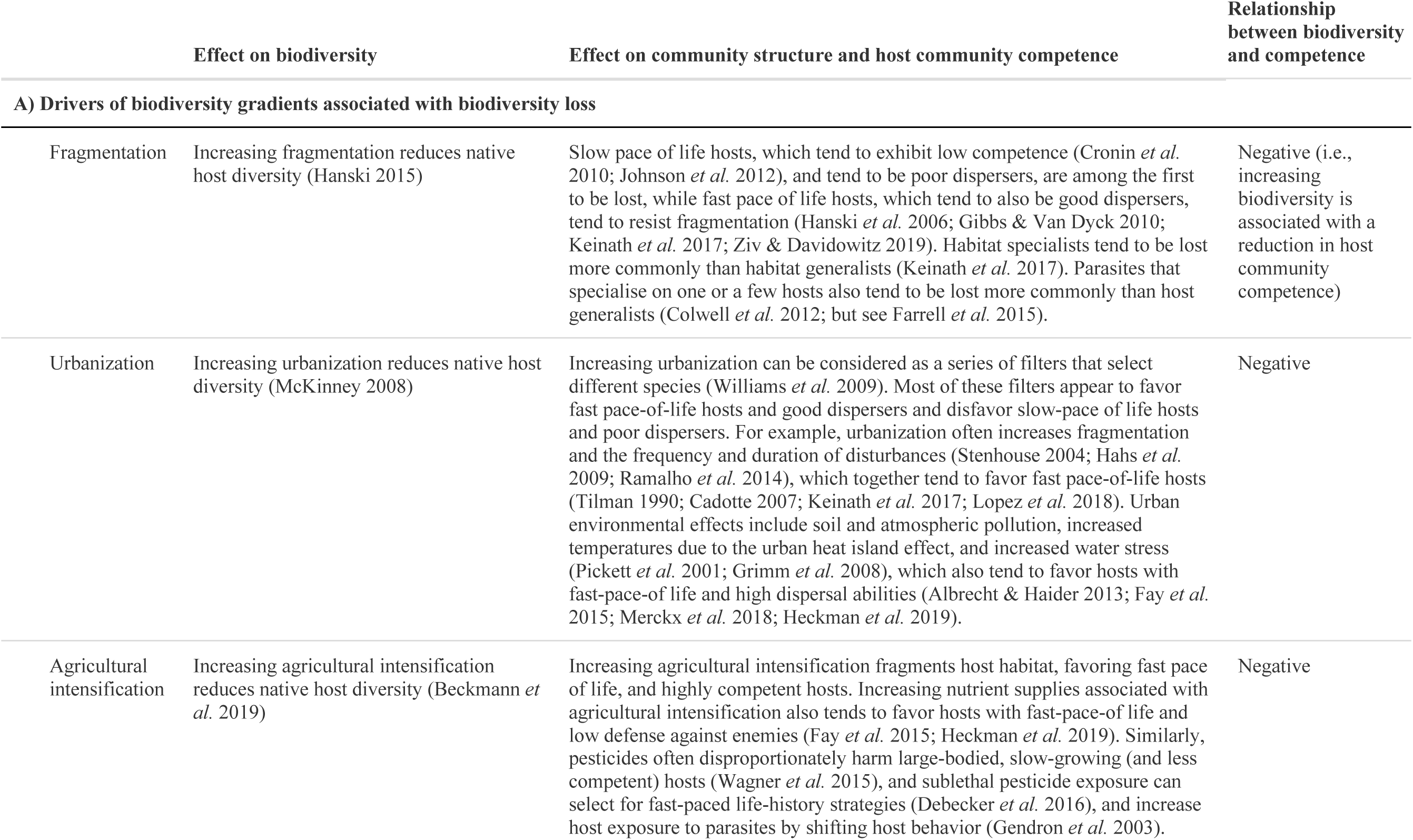
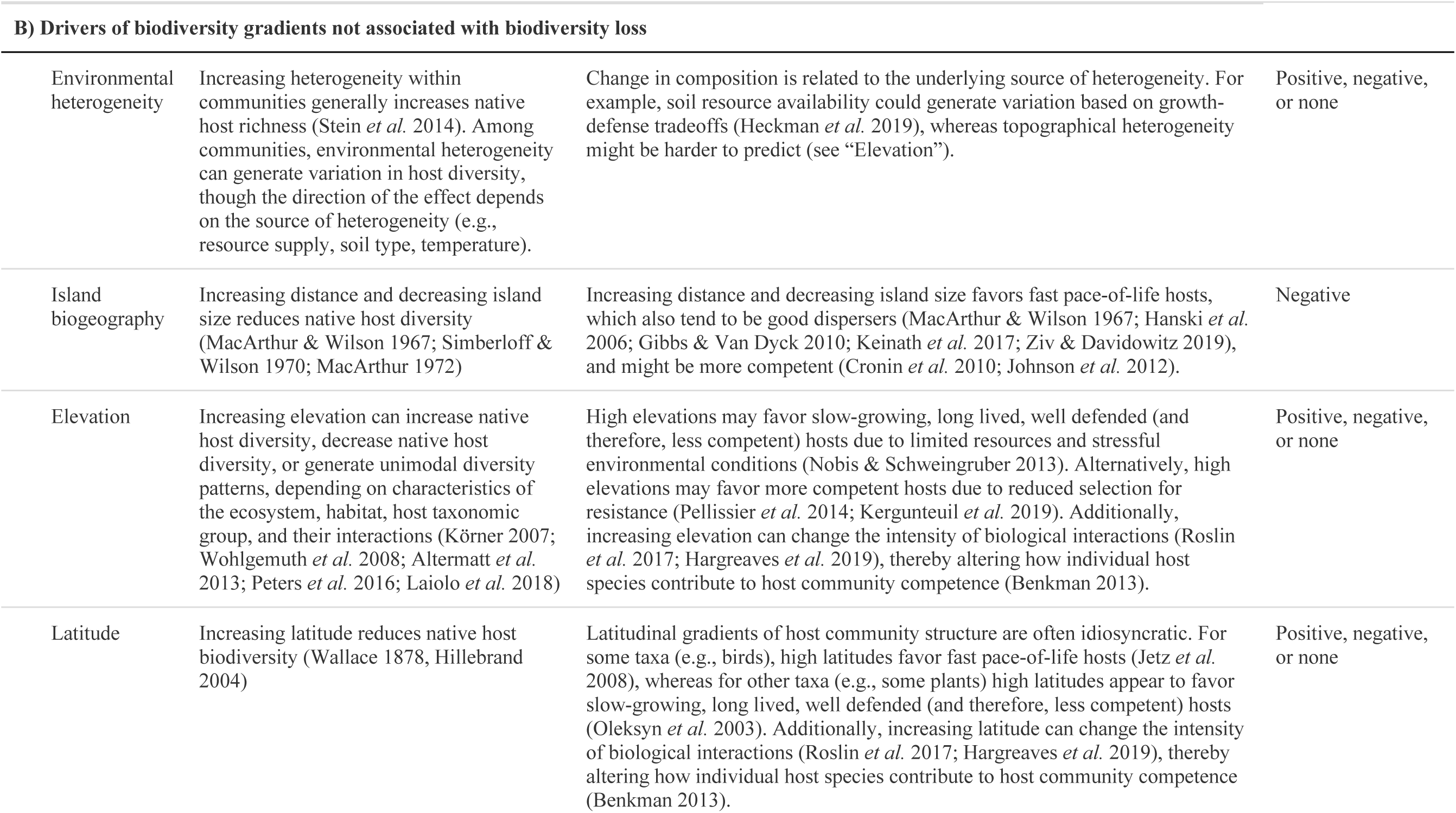
Common drivers of local biodiversity loss and expected impacts on host community structure

While many studies focus on measuring the diversity of host species in the context of disease, the structure of host communities can also be measured in the context of disease using characteristics of host species or host functional traits (Johnson *et al.* 2013; Halliday *et al.* 2019; Kirk *et al.* 2019), resulting in trait-based measures of host community competence. This approach, which has rapidly gained traction in disease ecology, suggests that host species that are the best able to spread diseases (i.e., the most competent hosts), often share particular suites of physiological traits (Huang *et al.* 2013; Martin *et al.* 2019; Becker & Han 2020). Thus, host community competence can be linked to distributions of important host traits across host communities (Johnson *et al.* 2015b; Liu *et al.* 2017). Importantly, several recent studies indicate that host community competence often covaries with host diversity, obscuring the true effect of host diversity, *per se*, on infectious disease risk (Johnson *et al.* 2015a; Young *et al.* 2017; Halliday *et al.* 2019). This covariance in host community competence and host diversity might, in turn, be driven by community disassembly or recolonization associated with biodiversity loss (Johnson *et al.* 2019; Rohr *et al.* 2020).

Biodiversity loss can drive the dilution effect because the most competent hosts also tend to be the species that remain or recolonize following biodiversity loss (Table 1a). One explanation for this pattern relates to host life history (Ostfeld & Keesing 2000; Previtali *et al.* 2012). Specifically, hosts with life history strategies that favor growth, reproduction, and dispersal, over defense against parasites (e.g., hosts exhibiting a fast pace of life), often contribute the most to disease in the communities that they occupy (i.e., act as disease amplifiers; Cronin *et al.* 2010; Johnson *et al.* 2012; Sears *et al.* 2015). Similarly, in a study of 2,277 vertebrate host species and 66 parasites, the best reservoir hosts (those with high abundance and diversity of parasites) were hosts with broad geographic ranges that invest heavily in reproduction and growth (Han *et al.* 2015b) (see also Luis *et al.* 2013). These fast pace-of-life hosts are also often the most resistant hosts to extinction (Hanski *et al.* 2006; Gibbs & Van Dyck 2010; Albrecht & Haider 2013; Fay *et al.* 2015; Keinath *et al.* 2017; Merckx *et al.* 2018; Ziv & Davidowitz 2019). Consequently, as host communities become fragmented or disturbed and biodiversity is lost, these fast pace-of-life, amplifying hosts remain, while their slow pace-of-life counterparts are lost (Joseph *et al.* 2013; Mihaljevic *et al.* 2014; Johnson *et al.* 2015a), leading to covariance between host diversity and host community competence. This hypothesis has been borne out for amphibian (Johnson *et al.* 2013), mammal (Ostfeld & LoGiudice 2003), and plant hosts (Liu *et al.* 2018). In two recent experiments, one using amphibian hosts (Johnson *et al.* 2019) and the other focused on plant hosts (Liu *et al.* 2018), dilution was not observed in communities that were disassembled randomly, but when communities disassembled naturally, biodiversity significantly reduced disease, lending further support to this hypothesis. Consequently, theory suggests that biodiversity gradients associated with biodiversity loss should result in dilution effects.

Whereas biodiversity loss is often linked to increased host community competence during community disassembly, the relationship between natural biodiversity gradients and host community competence is less clearly defined (Table 1b). For example, increasing elevation can increase host diversity, decrease host diversity, or generate unimodal diversity patterns, depending on characteristics of the ecosystem, habitat, host taxonomic group, and their interactions (Körner 2007; Wohlgemuth *et al.* 2008; Altermatt *et al.* 2013; Peters *et al.* 2016; Laiolo *et al.* 2018). Similarly, increasing elevation can select for more poorly-defended hosts when there is reduced selection for resistance at high elevations (Pellissier *et al.* 2014; Kergunteuil *et al.* 2019), but might also favor slow-growing, long-lived, well-defended hosts due to limited resources and stressful environmental conditions at high elevation (Nobis & Schweingruber 2013). Consequently using host competence to predict biodiversity-disease relationships along elevational gradients is challenging.

Different drivers of biodiversity gradients might also influence whether and when contingencies arise in the strength and direction of biodiversity-disease relationships (e.g., Halliday & Rohr 2019). For example, it has been proposed that biodiversity-disease relationships should be strongest at local scales and in tropical regions, where biotic interactions are strongest, and should weaken as spatial scale and (absolute values of) latitude increase and the strength of biotic interactions declines (Wood & Lafferty 2013; Johnson *et al.* 2015a; Cohen *et al.* 2016; Halliday & Rohr 2019; Liu *et al.* 2020; Rohr *et al.* 2020) (but see Magnusson *et al.* 2020). This effect might be particularly strong among studies that depend on biodiversity loss if biodiversity loss generates consistent patterns of host community competence, and might be weaker or even reverse among studies that do not depend on biodiversity loss depending on the relationship between biodiversity and host community competence (Table 1). Thus, moderation of the dilution effect might differ among studies that do not involve biodiversity loss and among studies that do.

In this study, we test whether the diluting effect of host diversity on disease risk varies between natural biodiversity gradients and biodiversity gradients that are associated with recent loss of native host species. We test this by reanalyzing four previously published meta-analyses that came to contradictory conclusions regarding generality in the dilution effect. Re-analyzing these data in the context of biodiversity loss reveals a unifying pattern: dilution effects are inconsistently observed for biodiversity gradients that are not associated with the loss of biodiversity (e.g., latitudinal, elevation, and habitat size gradients, or environmental heterogeneity), but are very regularly observed for biodiversity gradients that are generated by disturbances that cause losses of native biodiversity (Table 1). These patterns are robust to misclassification of as many as 50% of the biodiversity gradients in these two categories. Incorporating biodiversity loss into tests of generality in the dilution effect further helps to unify understanding of contingencies in the biodiversity-disease relationships, suggesting that scale-dependency should weaken the dilution effect when biodiversity gradients do not involve biodiversity loss, but may strengthen the dilution effect when biodiversity gradients are driven by biodiversity loss. Together, these results help to resolve one of the most contentious issues in disease ecology: the generality of the dilution effect.

## Methods

### Does biodiversity loss underlie the dilution effect of biodiversity?

To test whether biodiversity loss can explain generality in the relationship between biodiversity and disease risk, we reanalyzed four previously-published meta-analyses. These four previously published studies used different selection criteria and modeling frameworks, focused on different subsets of host and parasite taxa, and came to different conclusions regarding the generality of the dilution effect (Table 2). Conclusions from these published syntheses were contradictory, suggesting that the dilution effect can be robust (Civitello *et al.* 2015; Magnusson *et al.* 2020), scale dependent (Halliday & Rohr 2019), or dependent on latitude, habitat, and parasite life history (Liu *et al.* 2020).

**Table 2.** Summary of key data syntheses studying generality in the relationship between biodiversity and disease risk

|  | Data types | Studies (unique) | Manuscripts (unique) | Moderators | Contingencies identified | Moderators that interact with biodiversity loss |
| --- | --- | --- | --- | --- | --- | --- |
| **Civitello et al. 2015** | All studies | 208 (123) | 45 (21) | Parasite type, lifecycle, functional group, specialization; Study type | None | Parasite type |
| **Haliday & Rohr 2019** | Studies with more than three unique diversity measures | 217 (48) | 37 (6) | Spatial scale | Spatial scale | Spatial scale |
| **Magnusson et al. 2020** | Observational studies | 120 (16) | 37 (9) | Spatial scale; Latitude; Geographic region | Stronger relationships in temperate regions | Spatial scale |
| **Liu et al. 2020** | Studies of non-agricultural plant communities | 136 (58) | 20 (13) | Parasite life history, symptom; Ecosystem type; Study design; Latitude | Ecosystem type; Study design; Parasite life history; Latitude | Parasite life history, symptom; Latitude |

The following manuscripts were not included because the underlying source of the biodiversity gradient could not be identified: J. N. Mills. Archives of Virology, 45-57 (2005); A.T. Strauss, et al. Ecol Monogr 86(4):393-411, (2016); Zimmermann et al. Acta Parasitologica 62: 493-501 (2017); J. A. Lau, S. Y. Strauss. Ecology 86, 2990-2997 (2005). Sin Nombre Virus from unpublished data in D. J. Salkeld et. al. Ecology Letters 16, 679-686 (2013).

We obtained data and code (when available) from these four publications. For each study in each dataset, we assigned the driver of the underlying biodiversity gradient, and whether or not that driver was associated with biodiversity loss (presented in Table 1) by reading the abstract and methods of each study. We could not identify the driver of biodiversity gradients in five studies (Table 2), so those studies were omitted from our analysis.

So that all four datasets could be analyzed using the same analytical approach, we transformed Spearman Rank correlations from Halliday & Rohr (2019) into Fisher’s Z following the methods provided in Liu et al (2020). Briefly, the Spearman rank correlation from each study was transformed into Fisher’s Z using the following equation: 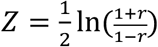, and the variance of Fisher’s Z was defined as 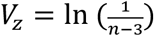. One study included a Spearman Rank correlation of -1. We therefore subtracted 1e^−5^ from the Spearman Rank correlation when calculating Fisher’s Z. We reconstructed the analyses performed in Civitello et al. (2015), Magnusson et al. (2020), and Liu et al. (2020), and analyzed Fisher’s Z from Halliday & Rohr (2019). Each model included whether or not the biodiversity gradient was associated with biodiversity loss as a moderator, and manuscript and parasite as random effects. All analyses were then performed using the R package metafor (Viechtbauer 2010).

### Are biodiversity-disease patterns robust to misclassification and whether or not studies included manipulative experiments?

We acknowledge that our classification of biodiversity gradients as being associated with biodiversity loss or not might be imprecise. For example, Rendón-Franco et al. (2014) measured diseases of small mammals in three different vegetation types: short grassland, tall grassland, and mesquite shrub, with the aim of acquiring a gradient of host richness and diversity. The factors that determined these three different vegetation types was unclear from the manuscript, so we assigned the driver of this biodiversity gradient as environmental heterogeneity, which is not associated with biodiversity loss (Table 1). However, it is equally possible that these three vegetation types were a reflection of different land-use histories, which would be associated with biodiversity loss, and that we therefore misclassified the underlying biodiversity gradient in this study.

To test whether our results were sensitive to misclassification in how we assigned drivers of biodiversity gradients, we randomly selected a proportion of studies, then randomly assigned the driver of biodiversity gradients in those studies, and re-analyzed the data, permuting this misclassification analysis 200 times for each misclassification rate.

In addition to problems of misclassification, assigning the underlying driver of biodiversity gradients in experiments can be problematic. Most experimental designs involve some kind of biodiversity loss; however, whether that loss is a random artifact of experimental design or represents a realistic example of biodiversity loss in nature depends on experimental design. Consequently, the relationship between biodiversity and disease risk in manipulative experiments is often sensitive to host composition (Venesky *et al.* 2014; Han *et al.* 2015a; Halliday *et al.* 2017). To our knowledge, only two studies have compared random and realistic biodiversity loss experimentally, with both studies finding that realistic biodiversity loss produced the strongest and most consistent dilution effects (Liu *et al.* 2018; Johnson *et al.* 2019). We therefore next dropped experiments from all datasets and re-analyzed the data.

### Does accounting for biodiversity loss explain inconsistencies among different data syntheses?

Finally, using the databasets, but excluding experiments, we tested whether inconsistencies among studies in the factors that modify the dilution effect could be explained by biodiversity loss. To this end, we re-analyzed the data, using the moderators tested in each original meta-analysis and an interaction between that moderator and whether or not the biodiversity gradient was associated with biodiversity loss.

## Results

Our reanalysis of the four previously published datasets revealed that biodiversity gradients associated with biodiversity loss consistently generated dilution effects (*p* < 0.01 for all studies), whereas other biodiversity gradients inconsistently generated dilution effects (Civitello et al.: *p* = 0.036; other studies: *p* > 0.05; Fig. 1). These patterns were robust to misclassification of the underlying source of biodiversity gradients in as many as 50% of the studies (Fig. 2). Moreover, the patterns were often robust to the exclusion of experimental studies, which can often test contrived community compositions. After excluding experiments in the Civitello et al., Halliday and Rohr, and Magnusson et al. datasets, biodiversity gradients associated with biodiversity loss still consistently generated dilution effects (*p* < 0.0001; *p* = 0.042; *p* < 0.0001, respectively), whereas gradients not clearly associated with biodiversity loss still did not (*p* = 0.07; *p* = 0.12; *p* = 0.14, respectively; Fig. 3). The exception was the Liu database (Table 2; Liu et al. 2020), where there was no significant dilution effect after excluding experiments (Biodiversity loss: *p* = 0.12; No biodiversity loss: *p* = 0.28).

**Figure 1.**
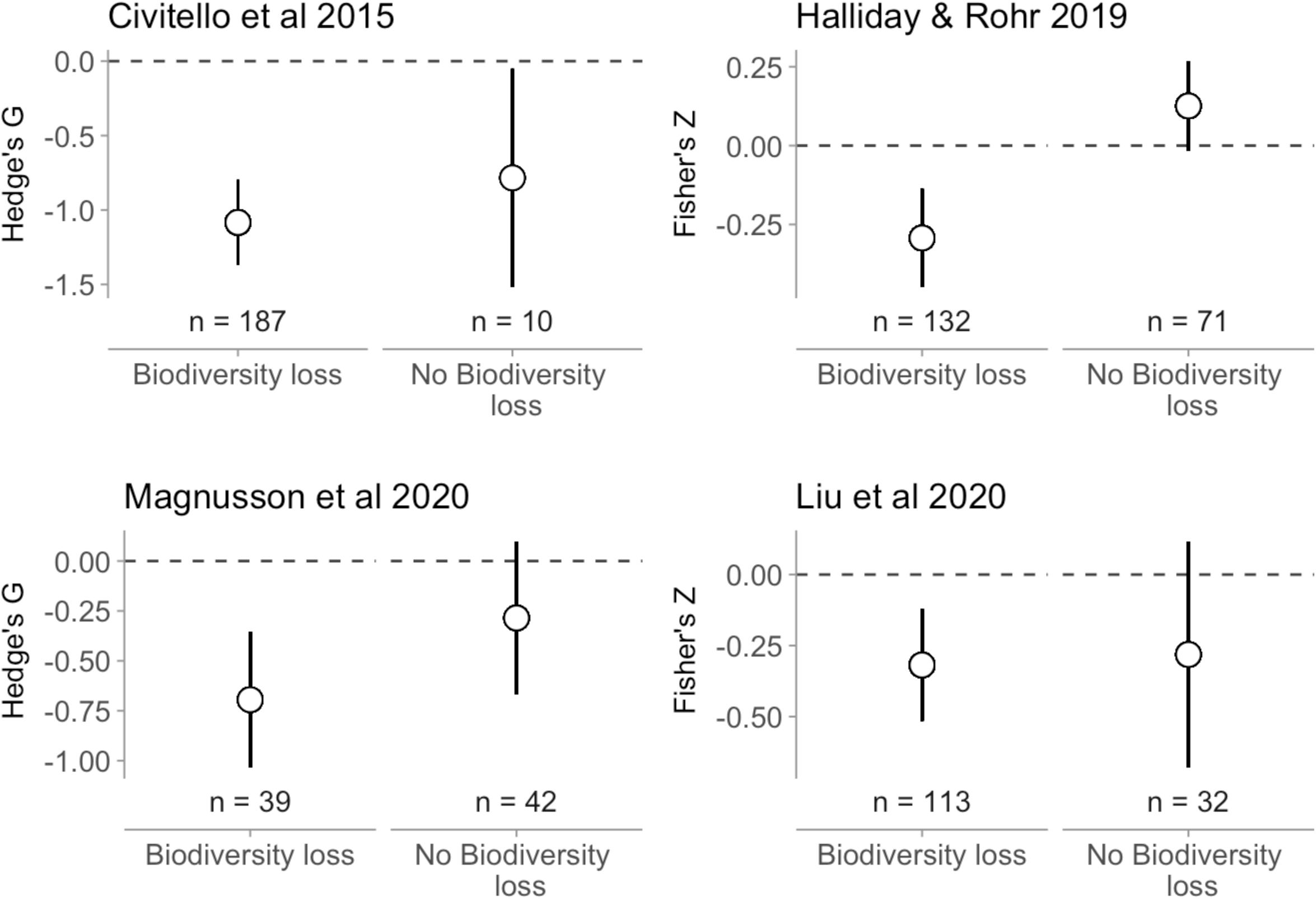
Effect of biodiversity loss on the dilution effect. Each panel corresponds to a separate meta-analysis of the dilution effect. The y-axis is a standardized effect size from the meta-analysis, aimed at estimating the strength of the dilution effect, with values below zero corresponding to a negative effect of biodiversity on disease risk (i.e., dilution). Points are model-estimated means, and error bars are model-estimated 95% confidence intervals. The dilution effect is robust across biodiversity gradients driven by biodiversity loss, but this effect is idiosyncratic across diversity gradients that do not involve biodiversity loss.

**Figure 2.**
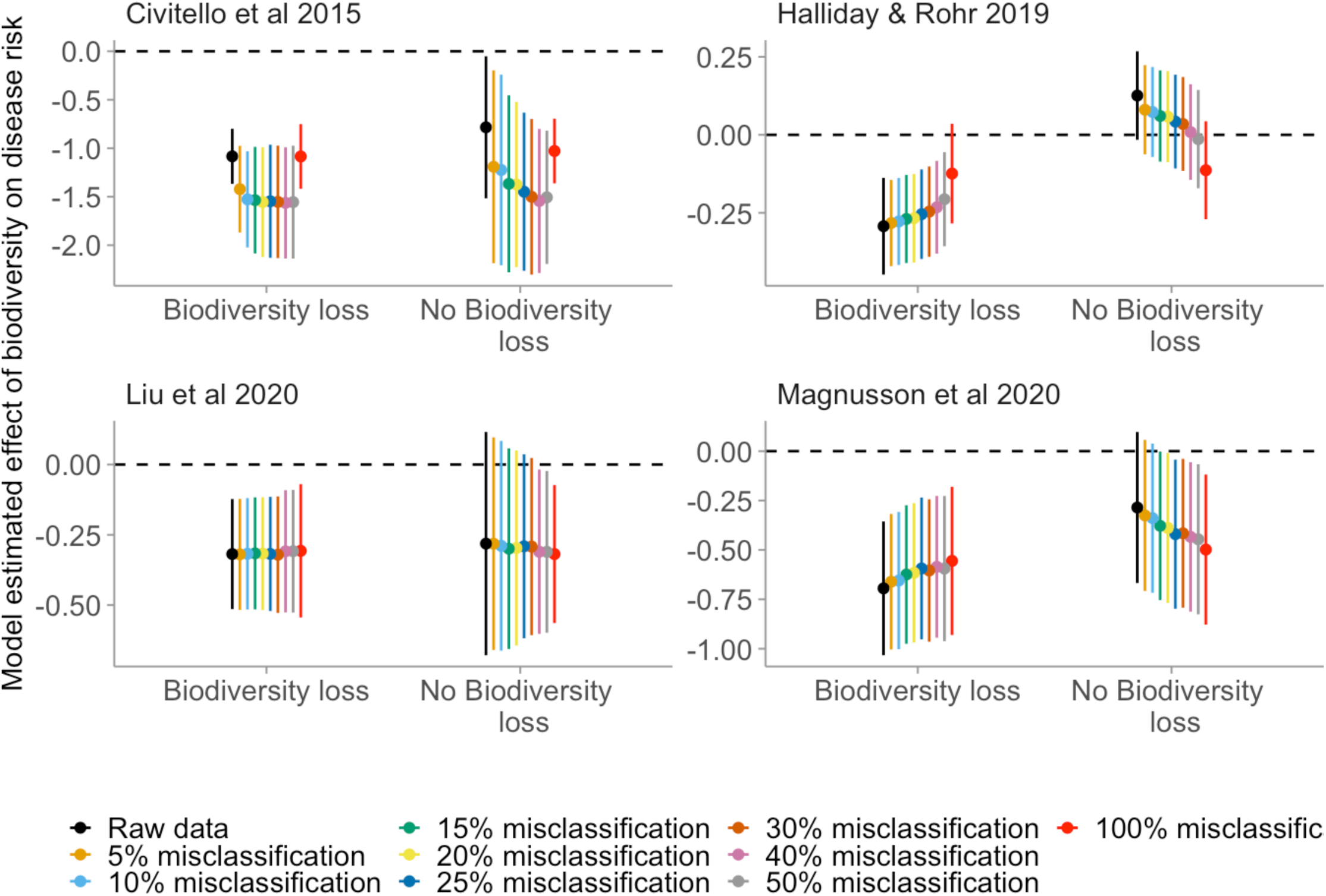
Effect of misclassification on moderation of the dilution effect by biodiversity loss. Each panel corresponds to a separate meta-analysis of the dilution effect. The y-axis is a standardized effect size from the meta-analysis, aimed at estimating the strength of the dilution effect, with values below zero corresponding to a negative effect of biodiversity on disease risk (i.e., dilution). Points are the average model-estimated mean, and error bars are he average model-estimated 95% confidence intervals across 200 simulations. The effect of biodiversity loss on the strength of the dilution effect is robus to misclassification of at least 10% and up to 50% of studies.

**Figure 3.**
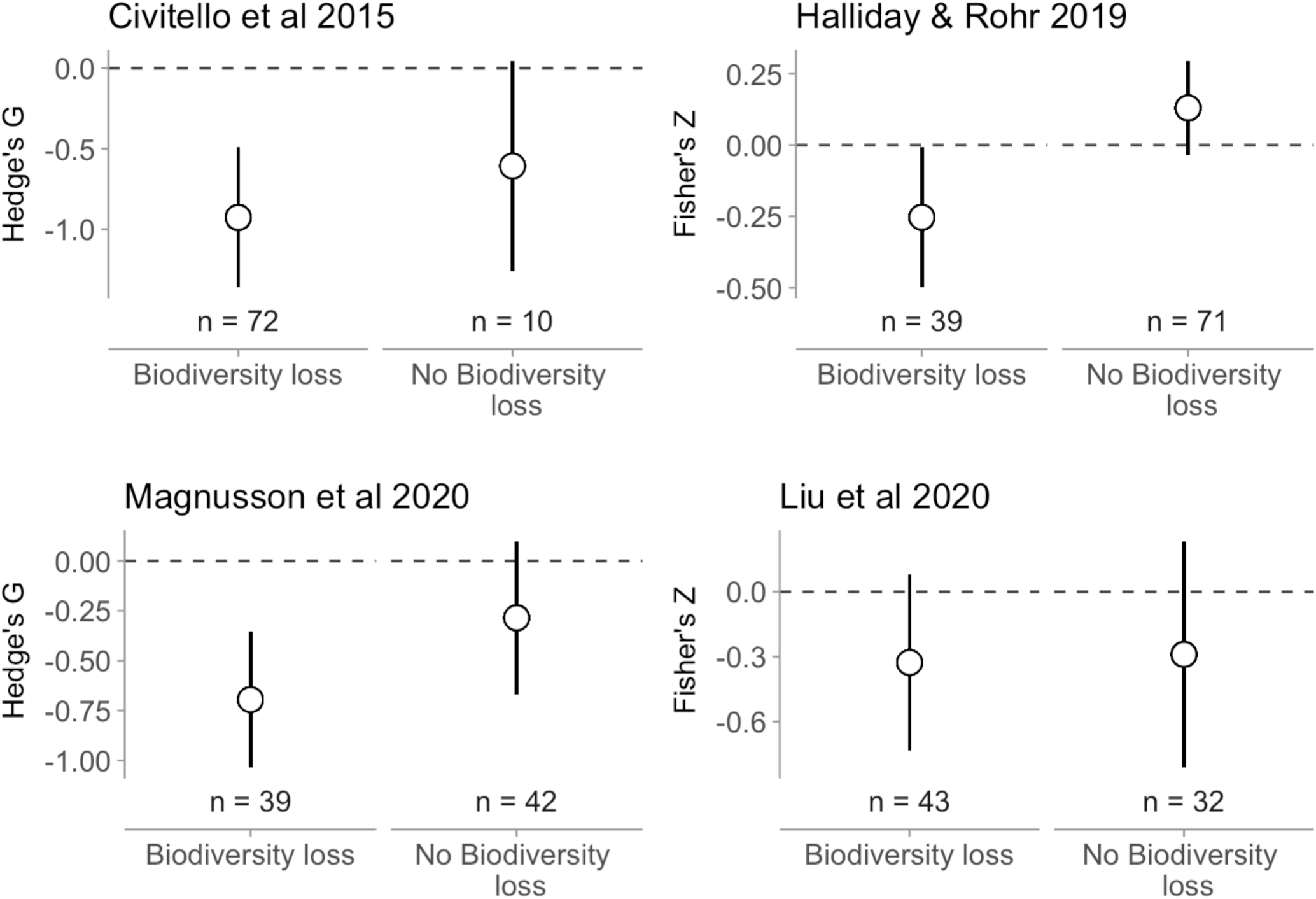
Effect of biodiversity loss on the dilution effect after excluding experiments. Panels correspond to different databases. Y-axes are standardized effect sizes, with values below zero corresponding to negative effects (i.e., dilution). Points are model-estimated means, and error bars are model-estimated 95% confidence intervals. With the exception of Liu, which was sensitive to study design, the dilution effect is robust across biodiversity gradients driven by biodiversity loss, but this effect is idiosyncratic across diversity gradients that do not involve biodiversity loss, even after excluding experiments.

Finally, we tested whether statistical interactions between biodiversity loss and moderators could explain inconsistencies among the four focal studies. The degree to which the four databases included gradients of biodiversity driven by biodiversity loss versus other factors resolved inconsistencies regarding spatial moderation of the dilution effect, but amplified descrepancies related to latitudinal gradients (Table 2; Fig. 4). Spatial scale significantly interacted with biodiversity loss in both studies that evaluated spatial scale (Halliday and Rohr: LRT 5.12, *p* = 0.024; Magnusson et al.: LRT 6.23, *p* = 0.013); the strength of dilution increased with scale for studies that involved biodiversity loss and weakened with scale for studies that did not (Fig. 4). In contrast to the consistency across studies in the scale patterns, we found a non-significant (LRT 2.40, *p* = 0.12) and significant (LRT 5.54, *p*=0.019) interaction between biodiversity loss and (absolute value of) latitude for the Magnusson et al. and Liu et al. datasets, respectively (Fig. 4). Moreover, the direction of these effects were opposite; in Liu et al., dilution weakened with increasing (absolute values of) latitude for biodiversity-loss studies, whereas in Magnusson et al., dilution strengthened with increasing (absolute values of) latitude for non-biodiversity-loss studies (Fig. 4).

**Figure 4.**
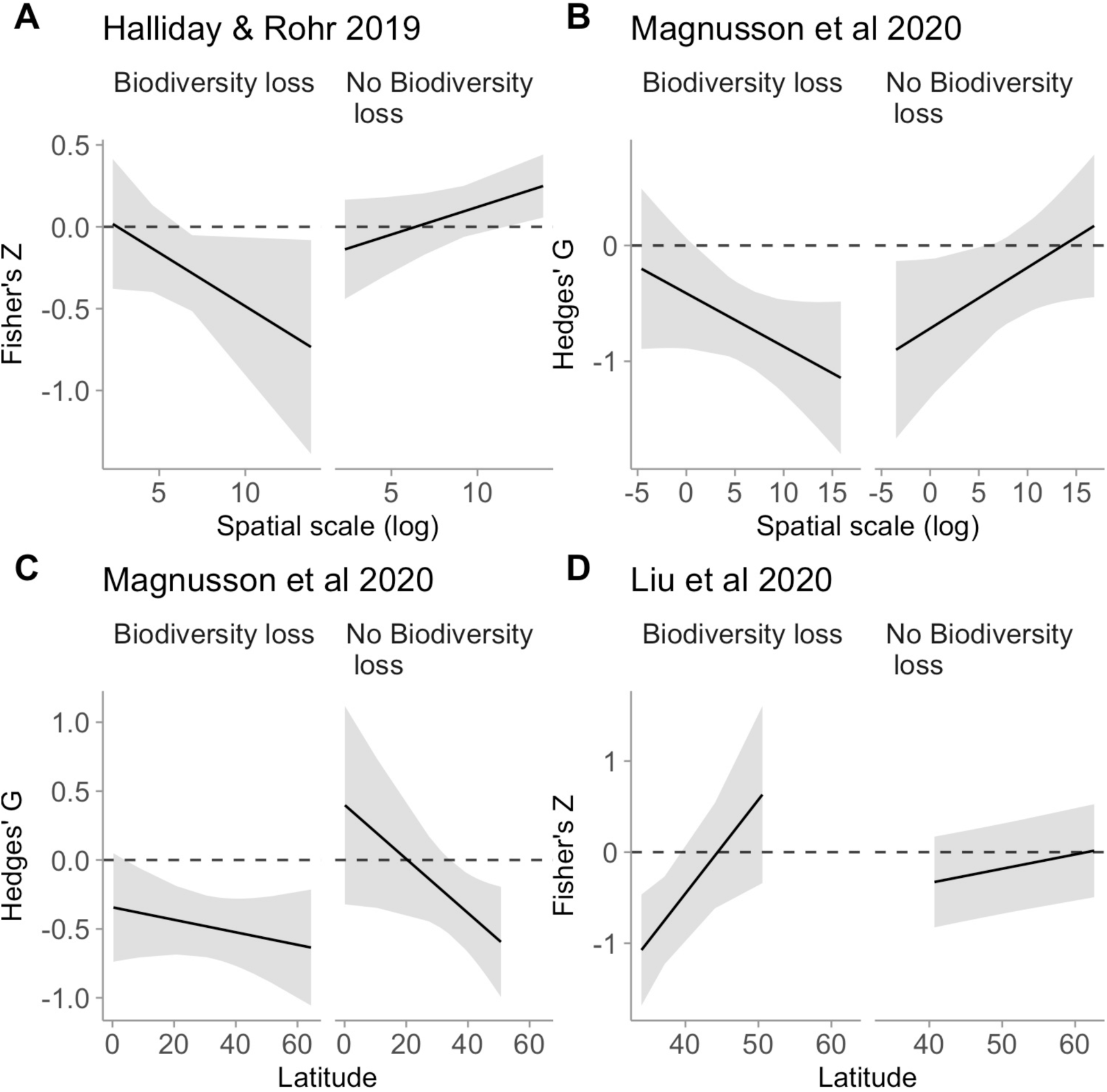
Effect of biodiversity loss on moderation of the dilution effect. Panels correspond to models of the interaction between biodiversity-loss and spatial scale (A & B) or latitude (C & D) for different meta-analyses, excluding experiments. The y-axis is a standardized effect size from the meta-analysis. Lines are model-estimated means, and ribbons are model-estimated 95% confidence intervals. Incorporating biodiversity loss resolves inconsistences in the effect of spatial scale, but not latitude.

## Discussion

This study shows broad evidence that biodiversity loss underlies the dilution effect. The effect of biodiversity loss on the dilution effect was robust to misclassification and whether or not studies included manipulative experiments. Furthermore, accounting for biodiversity loss explained some inconsistencies among prior data syntheses. Together, these results provide important context for understanding the role that native biodiversity plays in protecting human wellbeing and ecosystem health, suggesting that preventing biodiversity loss can proactively reduce infectious disease risk (Rohr *et al.* 2020).

Because community disassembly often favors more competent hosts (Table 1), we expected that biodiversity loss would commonly result in dilution effects. Our reanalysis of four published datasets is consistent with this idea: dilution effects were commonly observed among biodiversity-loss studies across all four datasets. However, we did not directly test whether dilution effects arise due to an increase in competent hosts, because most published studies do not report the identities or abundances of (potentially) diluting host species. Future studies should test for generality in this mechanism directly by comparing host community structure (including traits associated with host community competence and host biodiversity) across a variety of biodiversity drivers (e.g., Halliday *et al.* 2019), and in a variety of study systems.

Because the relationship between host community competence and biodiversity is often unpredictable along natural biodiversity gradients (Table 1), we expected that gradients not associated with biodiversity loss would inconsistently result in dilution effects. Our results support this idea: dilution effects were inconsistently observed among non-biodiversity-loss studies in three out of four datasets. However, these results do not suggest that dilution effects only occur when biodiversity gradients are associated with biodiversity loss. Importantly, even when there is no net association between host diversity and community competence, increasing biodiversity can still reduce disease risk of parasites that are specialized to infect a small number of host species by modulating host density (i.e., via encounter reduction; Mitchell *et al.* 2002; Keesing *et al.* 2006). Encounter reduction, in turn, might be particularly relevant when gradients include seasonality (e.g., latitude, elevation) that affects peak prevalence and the duration of the epidemic season. Thus, biodiversity gradients that are not associated with biodiversity loss could still generate consistent dilution effects via encounter reduction for specialist parasites. Understanding the degree to which biodiversity influences disease risk among specialists versus generalists in the context of biodiversity loss therefore remains an important topic for future studies.

Our prediction that biodiversity loss underlies the dilution effect was grounded in host community competence, because host communities become more competent as biodiversity is lost (e.g., Johnson *et al.* 2013; Liu *et al.* 2017); however, biodiversity loss could also influence the dilution effect by other potential mechanisms. As an example, biodiversity loss does not necessarily alter nutrient availability, but nutrient availability can underly a natural biodiversity gradient, with implications for higher trophic levels (Grace *et al.* 2016; Cappelli *et al.* 2019). Gradients that are or are not associated biodiversity loss could also differ in host abundance or density, connectivity of hosts and parasites, or host temporal turnover (Keesing *et al.* 2006, 2010; Young *et al.* 2014; Johnson *et al.* 2015a).

Our results also suggest that statistical interactions between biodiversity loss and spatial scale might be sufficient to explain inconsistencies among the four focal studies, but that interactions between biodiversity loss and latitude are not. However, as in prior studies on the dilution effect, we wish to emphasize that our analysis of spatial scale might be sensitive to the scarcity of studies conducted at the largest spatial scale and to a variety of study characteristics linked to spatial scale, including the metrics used to estimate diversity and disease, study design, and parasite type (Halliday & Rohr 2019). Importantly, both datasets that tested spatial scale only included one global study where the underlying gradient involved biodiversity loss (Derne *et al.* 2011) and only one global study where the underlying gradient did not involve biodiversity loss (Wood *et al.* 2017). Consequently, we cannot rule out the possibility that these results could change if future studies filled these research gaps. Nevertheless, incorporating biodiversity loss resolved inconsistencies among studies related to spatial moderation of the dilution effect.

Even among datasets where biodiversity loss interacted with scale or latitude, the direction and magnitude of these interactions was not always consistent with theory. Specifically, theory predicts that increasing spatial scale and (absolute values of) latitude should weaken the dilution effect, because biotic interactions tend to weaken with increasing spatial scale and (absolute values of) latitude (Wood & Lafferty 2013; Johnson *et al.* 2015a; Cohen *et al.* 2016; Halliday & Rohr 2019; Liu *et al.* 2020; Rohr *et al.* 2020) (but see Magnusson *et al.* 2020). We therefore expected that if host community competence drives the dilution effect (Johnson *et al.* 2013), and this process occurs more commonly when native biodiversity is lost (Table 1), then this moderating effect of latitude and spatial scale would be strongest among biodiversity-loss studies. Consistent with this hypothesis, increasing latitude weakened the dilution effect in biodiversity-loss studies, though this effect was only observed in one dataset (Liu *et al.* 2020). In contrast, increasing scale increased the strength of the dilution effect among biodiversity-loss studies. We suggest that this result might be more statistical than biological: among non-biodiversity-loss studies where biodiversity is not associated with host community competence, large spatial scales can confound biodiversity gradients with changes in species pools, weakening dilution effects (Wood & Lafferty 2013; Rohr *et al.* 2020). In contrast, among biodiversity-loss studies where biodiversity is associated with community competence regardless of the underlying species pool, increasing scale could strengthen the dilution effect, particularly if large-scale studies capture a larger portion of the biodiversity gradient than smaller-scale studies, and their biodiversity-disease relationships favor dilution over the majority of the gradient (i.e., they are right-skewed; Halliday & Rohr 2019; Rohr *et al.* 2020). These results highlight the need for studies that measure biodiversity gradients across spatial scales to better disentangle conditions under which spatial scale and latitude moderate the dilution effect.

Together, the results of this study highlight the need to consider drivers of biodiversity gradients when predicting the role of biodiversity in influencing infectious disease. Specifically, our results suggest that dilution effects may occur less commonly for biodiversity gradients that are not associated with the loss of biodiversity, but occur regularly for biodiversity gradients that are generated by disturbances that cause losses of native biodiversity. These results are consistent with a growing body of literature suggesting that the role of biodiversity in regulating ecosystem processes depends on characteristics of species or individuals present in those ecosystems (Mouillot *et al.* 2011; Allan *et al.* 2015; Leitão *et al.* 2016; Van de Peer *et al.* 2018; Bagousse-Pinguet *et al.* 2019; Start & Gilbert 2019; Heilpern *et al.* 2020). These results therefore provide clarity in an increasingly polarized debate. Specifically, because characteristics of host communities often predictably change with biodiversity loss, these results suggest that biodiversity loss generally exacerbates infectious disease risk.

## Acknowledgements

We are grateful for insightful suggestions from D. Civitello, M. Jalo, and members of the Laine Lab. This work was supported by the University of Zürich and by grants from the Academy of Finland (296686) to A-LL and the European Research Council (Consolidator Grant RESISTANCE 724508) to A-LL.

