## Supplement for "Biodiversity loss underlies the dilution effect of biodiversity"

**Supplementary material**

Incorporating biodiversity loss into previously published tests of moderation of the dilution effect.

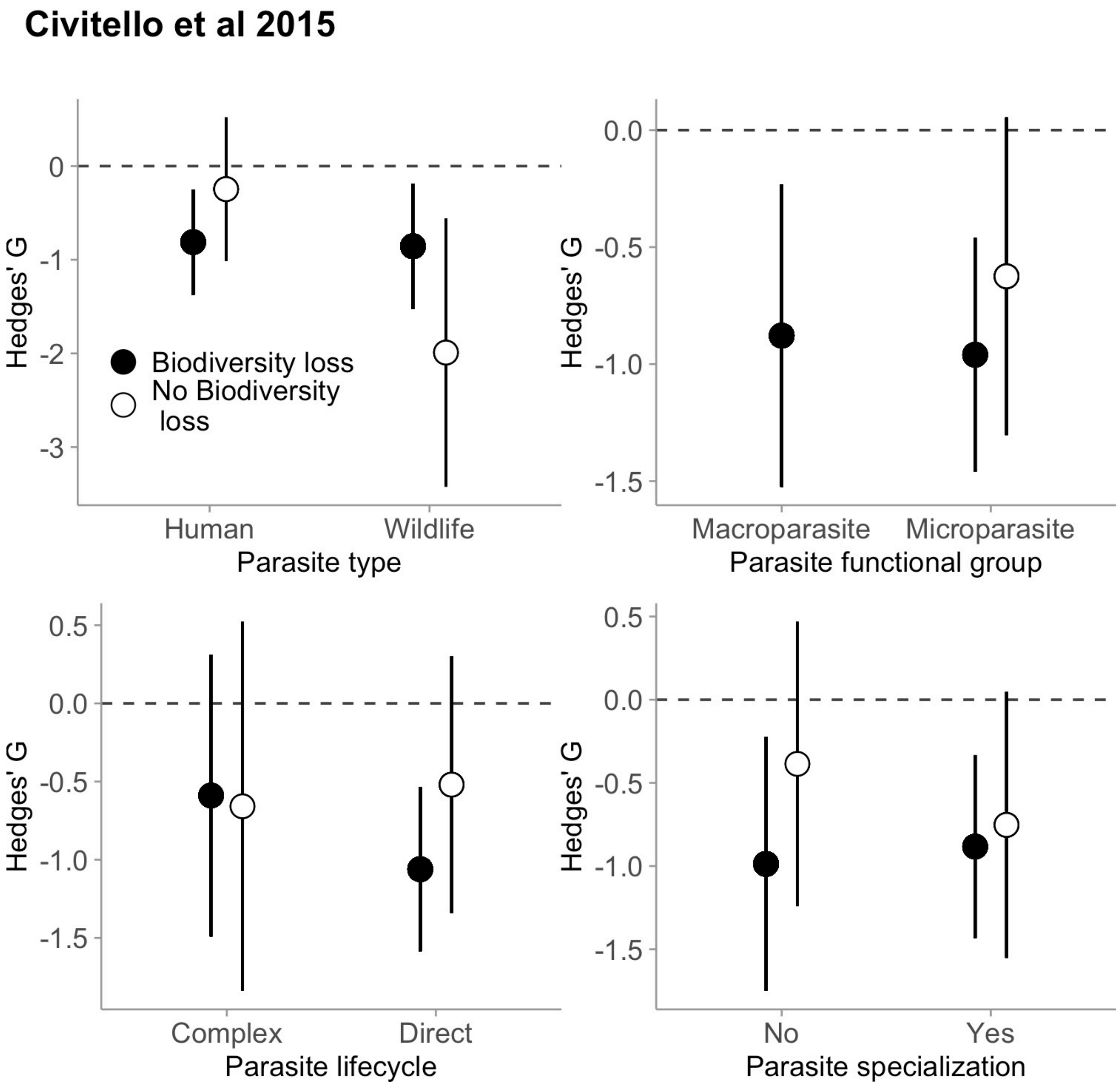


**Figure S1.** Effect of biodiversity loss on moderation of the dilution effect using data from Civitello et al 2015, excluding experiments. Points are model-estimated means; error bars are model-estimated 95% confidence intervals.

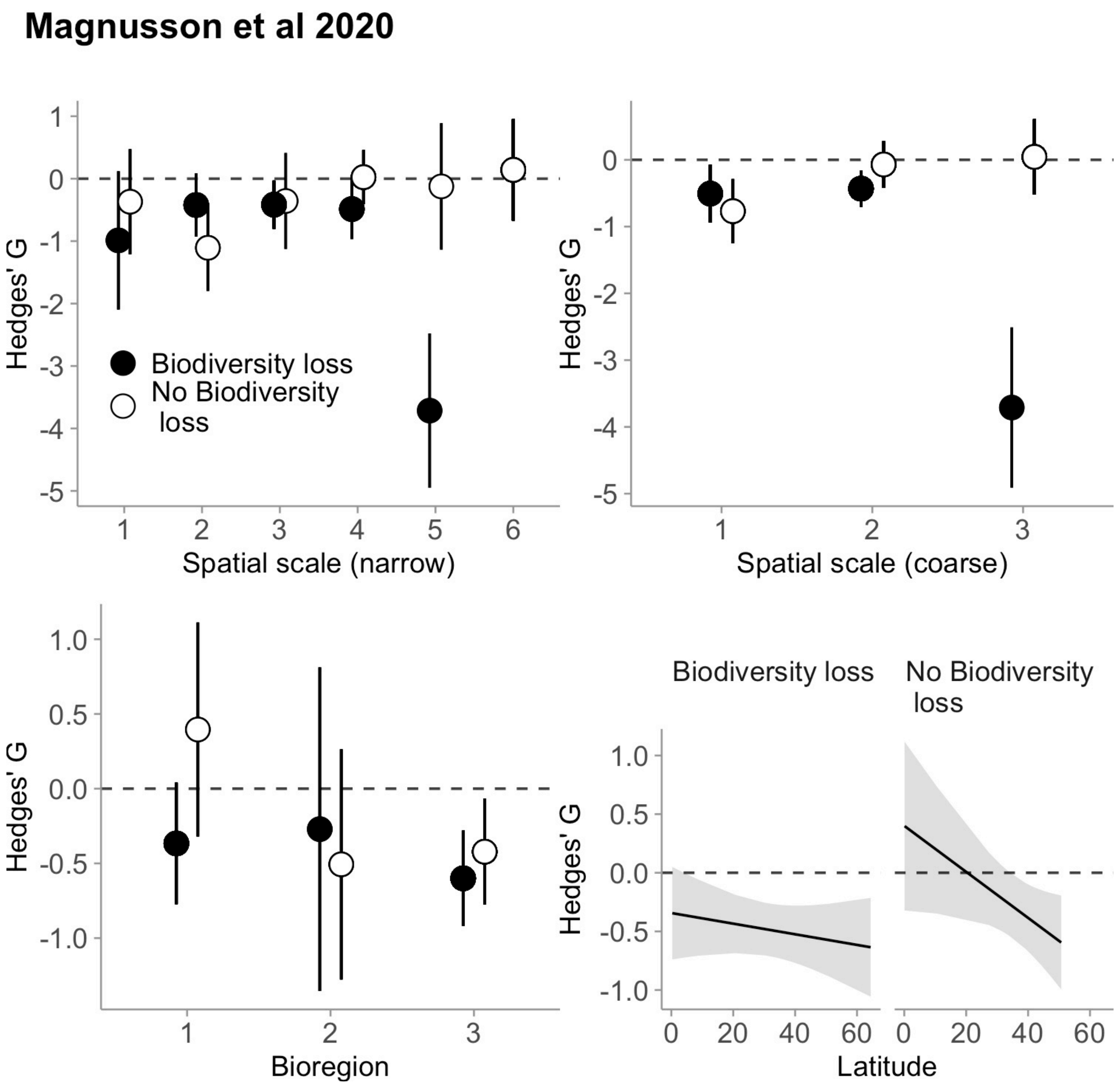


**Figure S2.** Effect of biodiversity loss on moderation of the dilution effect using data from Magnusson et al 2020, excluding experiments. Points and lines are model-estimated means; ribbons and error bars are model-estimated 95% confidence intervals.

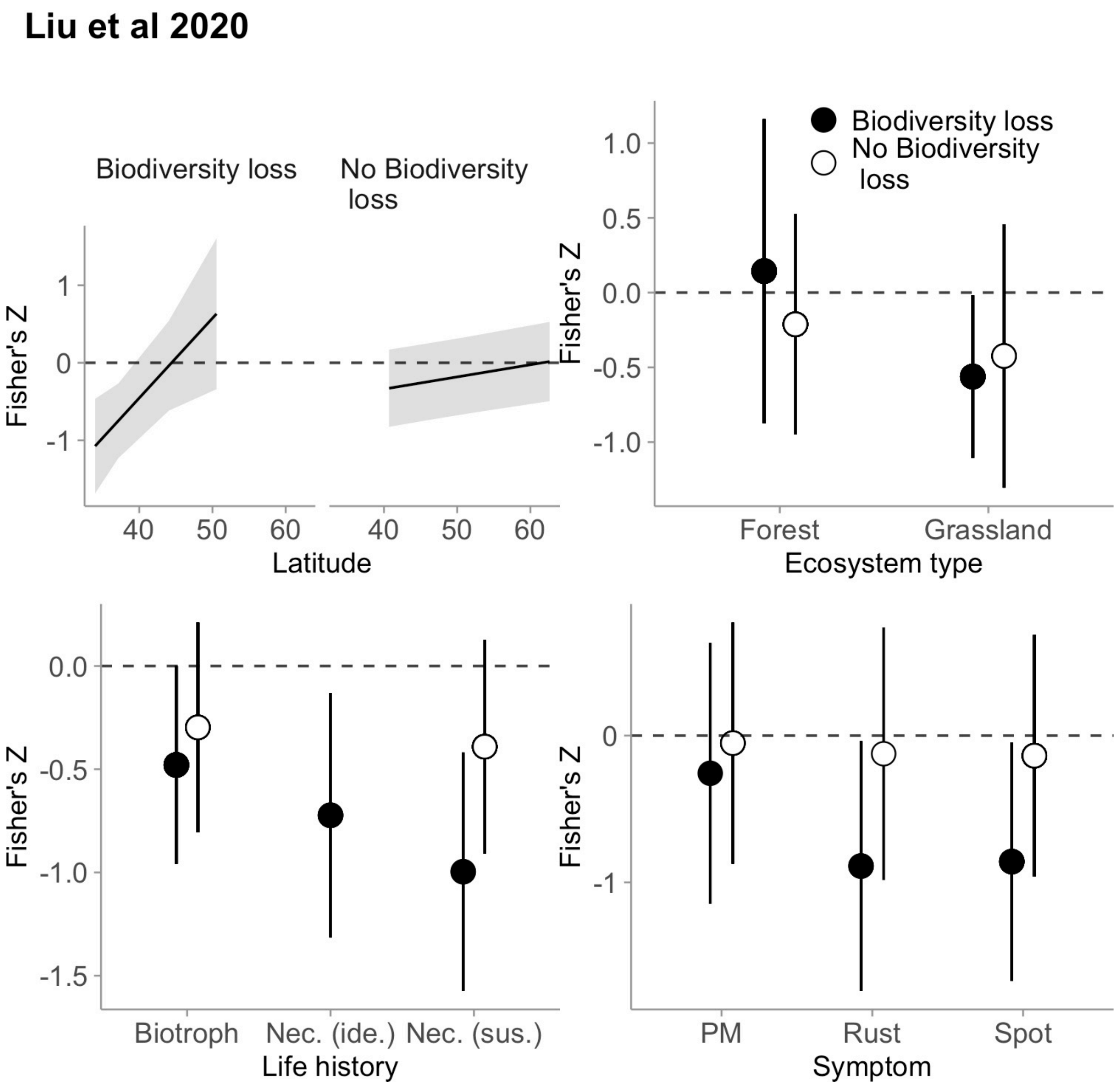


**Figure S3.** Effect of biodiversity loss on moderation of the dilution effect using data from Liu et al 2020, excluding experiments. Points and lines are model-estimated means; ribbons and error bars are model-estimated 95% confidence intervals.
